# Reservoir host community and vector density predict human tick-borne diseases across the United States

**DOI:** 10.1101/2020.10.15.341107

**Authors:** Michael B. Mahon, Jason R. Rohr

## Abstract

In the United States, tick-borne disease cases have tripled since the 1990s and cost upwards of 10 billion USD annually. Tick density and densities and diversity of non-human mammalian reservoir hosts are hypothesized to drive tick-borne disease dynamics and are targets for interventions. Here, we relate human prevalence of four tick-borne diseases (Lyme disease, monocytic ehrlichiosis, granulocytic anaplasmosis, and babesiosis) to tick and reservoir host community data collected by the U.S. National Ecological Observatory Network (NEON) across the contiguous U.S. We show that human disease prevalence is correlated positively with tick and reservoir host densities and negatively with mammalian diversity for Lyme disease and ehrlichiosis, but positively for anaplasmosis and babesiosis. Our results suggest that the efficacy of tick-borne disease interventions depends on tick and host densities and host diversity. Thus, policymakers and disease managers should consider these ecological contexts before implementing preventative measures.

**Significance:** Tick-borne disease incidence has increased in the United States over the last three decades. Because life-long symptoms can occur if reactive antibiotics are not administered soon after the tick bite, prevention is imperative. Yet, control of tick-borne zoonoses has been largely unsuccessful, at least partly because of a limited understanding of the ecological complexities of these diseases, especially non-Lyme disease tick-borne zoonoses. We use continental-scale data to quantify the relationships among four tick-borne diseases and tick and reservoir host communities, revealing that disease incidence is driven by a combination of tick densities and reservoir host densities and diversity. Thus, the efficacy of tick-borne disease interventions is likely dependent on these ecological contexts.

## Introduction

Vector-borne diseases and tick-borne diseases, specifically, are on the rise globally (1). In the United States, tick-borne disease incidence has more than doubled since 2004 and Lyme disease incidence has tripled since the 1990s (2, 3). There are an estimated 240,000-440,000 new cases of Lyme disease annually (4). Lyme disease costs the United States >$1 billion in healthcare costs (5), and $5 to $10 billion in economic and societal costs annually (6), and is only one of several major tick-borne disease in the U.S. Antibiotics are often ineffective at preventing life-long symptoms of bacterial tick-borne diseases (e.g. Lyme disease, human granulocytic anaplasmosis) if they are not prescribed soon after the tick bite (7) and, thus, effective prevention of tick-borne diseases is crucial. However, control of tick-borne zoonoses has been largely unsuccessful, in part because of a limited understanding of the ecological complexities and drivers of these diseases (1, 8, 9).

Tick densities, reservoir host densities, and reservoir host diversity are hypothesized drivers of tick-borne diseases (Fig. 1) (1, 10, 11). The causative agents of human Lyme disease (*Borrelia burgdorferi* [*sensu lato*]), human granulocytic anaplasmosis (*Anaplasma phagocytophilum*; hereafter anaplasmosis), and human babesiosis (*Babesia microti*) are transmitted by *Ixodes scapularis* (eastern blacklegged ticks) in the eastern U.S. and *Ixodes pacificus* (western blacklegged ticks) in the western U.S., while the causative agent of human monocytic ehrlichiosis (*Ehrlichia chaffeensis*; hereafter ehrlichiosis) is transmitted by *Amblyomma americanum* (lone star ticks) throughout the U.S. (12–14). Larval ticks become infected following a blood-meal from an infected host, such as mammals (e.g. rodents, insectivores, and scurids (15)) that can serve as reservoir hosts (organisms that maintain and transmit pathogen populations (16)) for these pathogens. Importantly, reservoir competence (ability for a host to transmit pathogens to uninfected vectors (16)) of mammalian host species varies among these pathogens (Fig. 1; Table S1) (15, 17, 18).

**Figure 1.**
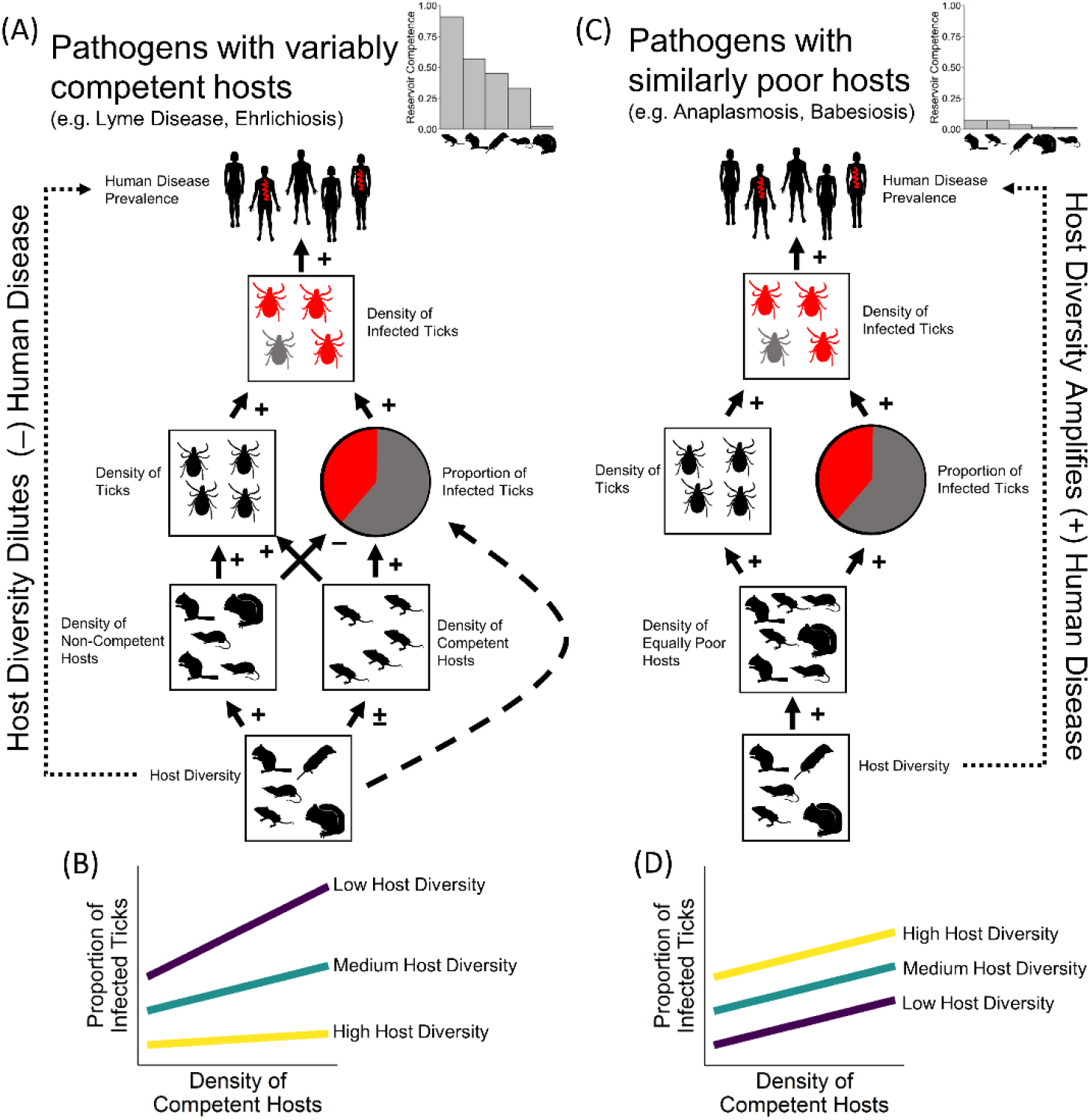
Conceptual diagram for linkages connecting wildlife to human prevalence of tick-borne diseases, per the dilution effect hypothesis. (A), hypothesized links among wildlife and human disease prevalence for pathogens with hosts that differ in reservoir competence (ability to transmit pathogen to uninfected ticks; coefficient of variation (CV) for tested *B. burgdorferi* reservoirs: 1.28, CV for tested *E. chaffeensis* reservoirs: 1.10). (B), hypothesized interaction between host diversity and competent host density, by which the effect of competent host density on tick infection prevalence is strongest when host diversity is low. (C), hypothesized links among wildlife and human disease prevalence for pathogens with hosts that are similarly poor reservoir hosts (CV for tested *A. phagocytophilum* [human strain] reservoirs: 0.80; CV for tested *B. microti* reservoirs: 0.81). (D), hypothesized additive relationship between host diversity and competent host density, such that the effect of increased host diversity in an area will increase reservoir density and, in turn, tick infection prevalence. In (A) and (C), links between density of infected ticks and human disease prevalence can be disrupted by human behavior (e.g. acaricides and avoidance). For each link in (A) and (C), signs are representative of whether relationship is positive (+), negative (−), or variable (±). Dotted arrows from host diversity to human disease prevalence in (A) and (C) are the overall, indirect effect of diversity on disease. Data for host reservoir competence in (A) and (C) are from ref (15, 17, 18).

Although there is some support for the hypotheses that densities of ticks and densities and diversity of wildlife hosts drive Lyme disease dynamics, most of the support is at local rather than country or continental scales (14, 15, 19) and has not incorporated human disease (e.g. reported cases) (10, 11, 19) (but see (20)). Linking wildlife and densities of infected ticks to human disease is difficult, because human behavior, such as avoidance and chemical deterrents (21), can disrupt this link. Further, support for the hypotheses that tick densities and wildlife host densities and diversity drive tick-borne diseases is lacking in systems other than Lyme disease. A major impediment to understanding the drivers of tick-borne diseases was a lack of broad-scale spatial datasets that combined reservoir host communities, tick densities, and tick infection prevalence with human disease incidence at similar scales (22). With the establishment of the U.S. National Ecological Observatory Network (NEON), there are now data on reservoir host communities, tick densities, and tick infection prevalence from across the U.S. collected using standardized methodologies that can be coupled with U.S. Center for Disease Control (CDC) data on human tick-borne disease incidence to finally elucidate the role of wildlife factors on human prevalence of tick-borne diseases.

The objectives of our study are to: 1) identify the combination of host and vector community variables that best explain human tick-borne disease prevalence across space and time, 2) evaluate the direct and indirect effects of these variables on human disease prevalence mediated through density of infected ticks, and 3) evaluate the human health burden of tick-borne diseases across a mammalian diversity gradient. In accordance with the dilution effect hypothesis (14, 23), we expect a negative biodiversity-disease relationship (dilution) when the most abundant hosts have the highest reservoir competence, because as rare host species are added to communities the mean competence of the community, and thus disease risk, decreases (assuming host community assembly is substitutive; i.e. competitive for niche space). We expect a negative biodiversity-disease relationship (amplification) when hosts do not or weakly differ in their reservoir competence because as rare hosts are added the mean competence of the host community would not change, whereas host densities and disease risk could increase if assembly is additive (Fig. 1; see Supplement Text for further details) (10, 14, 23). Hence, we predict that increasing small mammal richness will dilute *B. burgdorferi* (causative agent of Lyme disease) and *E. chaffeensis* (causative agent of ehrlichiosis), because the most abundant mammal hosts are typically the most competent for these pathogens (Fig. 1, Table S1), but we expect increasing small mammal richness to amplify *A. phagocytophilum* (causative agent of anaplasmosis) and *B. microti* (causative agent of babesiosis), because small mammal hosts exhibit similarly poor competencies for these pathogens (Fig. 1, Table S1).

To address these objectives, we linked tick density, and mammal density and diversity data collected by NEON from 2014-2018 to CDC data on human cases of Lyme disease, anaplasmosis, babesiosis, and ehrlichiosis gathered for the same counties and time as the NEON data (see Table S2 for specific county-year replicates included in analyses; Fig. S1). Because of differences in disease reporting and data availability (*Materials and Methods*), sample sizes (site-year replicates) for model selection were variable (*n* = 95 for Lyme disease, *n* = 116 each for anaplasmosis and ehrlichiosis, and *n* = 57 for babesiosis). To analyze correlations among ecological factors and human tick-borne disease prevalence, we used generalized linear mixed effects models with a binomial distribution and county as a random effect (24), and conducted model selection of all main effects and biologically relevant two-way interactions among wildlife variables (interactions between tick and host densities and host diversity and densities; see Table S3 for competing models). To account for differences in questing height of blacklegged ticks along a north-south gradient (25), we included latitude as a covariate for blacklegged tick-borne diseases. Additionally, to account for potential climate-related differences in relationships between wildlife variables and human disease incidence (20), all models included covariates of mean annual temperature and annual precipitation. The previously described analyses did not include tick infection data because these data were collected at only 13 NEON sites (*Materials and Methods*), which would have reduced the statistical power to test our hypotheses. To investigate direct and indirect effects of tick and host community metrics on human disease prevalence mediated through density of infected ticks, we used sequential regressions (see Fig. 1 for *a priori* relationships).

## Results

Tick density, reservoir host density, and small mammal species richness predicted human disease prevalence, but the direction of effects and interactions among these variables differed across diseases (Table 1, Table S3; Fig. 2). For Lyme disease and anaplasmosis, the relationship between reservoir host density and human disease prevalence became more positive as tick density increased (Fig. 2A,B). Thus, for Lyme disease and anaplasmosis, the reduction of a single tick per 1,000 m^2^ would reduce incidence by 29.4% and 26.1% at median host densities, and 99.9% and 102.0% at high host densities (4 mice per 10,000 m^2^ for Lyme diseaase and 7 small mammal individuals per 10,000 m^2^ for anaplasmosis), respectively. For babesiosis and ehrlichiosis, the reduction of tick density by a single individual per 1,000 m^2^ would reduce disease incidence by 27.1% and 25.8%, respectively. Like reducing ticks, decreasing reservoir host densities also reduces human tick-borne diseases (Fig. 2). In fact, when other factors in the model are at median values, a reduction in one tick per 1,000 m^2^ and a reduction of one reservoir host individual per 10,000 m^2^ (lowering of deer density category for ehrlichiosis) is predicted to prevent annually ~2,300 and ~2,500 total (across all four diseases) U.S. tick-borne disease cases, respectively, and ~300 and ~20 U.S. Lyme disease disability-adjusted life years (DALYs) (26), respectively, relative to 2017 data (Fig. S2). DALY information is unavailable for the other three tick-borne diseases, which should be addressed in future research given that tick-borne diseases vary in symptoms and virulence (2, 8, 26) and, thus, cases of tick-borne diseases do not represent human disease burden.

**Table 1.** Regression coefficients and statistics from best-fit models from Table S3. Predictor variables were blacklegged tick density, lone star tick density, *B. burgdorferi* reservoir density, small mammal richness, small mammal density, and white-tailed deer density. Models included a random effect of county. All models included covariates of annual mean temperature and annual precipitation. Lyme disease models included covariates of CDC reporting type. R^2^ values represent marginal/conditional (fixed/random + fixed) R^2^.

| Model/Variable | Estimate<br>(SE) | DF | Chisq | P |
| --- | --- | --- | --- | --- |
| Lyme disease ($w = 0.99$, $R^2 = 0.17/0.78$, $n = 95$, $X^2(5) = 38.323$, $p < 0.001$) | | | | |
| Blacklegged tick density | -0.177 (0.12) | 1 | 2.12 | 0.146 |
| *B. burgdorferi* reservoir abundance | 0.299 (0.21) | 1 | 2.09 | 0.148 |
| Small mammal richness | 0.057 (0.08) | 1 | 0.58 | 0.445 |
| Blacklegged tick density* *B. burgdorferi* reservoir density | 0.217 (0.05) | 1 | 19.99 | < 0.001 |
| *B. burgdorferi* reservoir density*Small mammal richness | -0.097 (0.03) | 1 | 12.25 | < 0.001 |
| Anaplasmosis ($w = 0.69$, $R^2 = 0.32/0.82$, $n = 116$, $X^2(4) = 17.346$, $p = 0.002$) | | | | |
| Blacklegged tick density | -0.946 (0.47) | 1 | 4.04 | 0.044 |
| Small mammal density | -0.065 (0.04) | 1 | 2.87 | 0.090 |
| Small mammal richness | 0.340 (0.12) | 1 | 7.44 | 0.006 |
| Black-legged tick density*Small mammal density | 0.236 (0.07) | 1 | 10.29 | 0.001 |
| Babesiosis ($w = 0.51$, $R^2 = 0.73/0.74$, $n = 57$, $X^2(4) = 20.938$, $p < 0.001$) | | | | |
| Blacklegged tick density | 0.240 (0.12) | 1 | 4.31 | 0.038 |
| Small mammal density | 0.875 (0.63) | 1 | 1.94 | 0.164 |
| Small mammal richness | 0.657 (0.55) | 1 | 1.42 | 0.234 |
| Small mammal density*Small mammal richness | -0.119 (0.06) | 1 | 3.36 | 0.067 |
| Ehrlichiosis ($w = 0.22$, $R^2 = 0.44/0.46$, $n = 116$, $X^2(3) = 19.61$, $p < 0.001$) | | | | |
| Lone star tick density | 0.230 (0.08) | 1 | 7.44 | 0.006 |
| Deer density | 1.660 (0.46) | 1 | 12.88 | <0.001 |
| Small mammal richness | -0.197 (0.11) | 1 | 3.37 | 0.066 |

**Figure 2.**
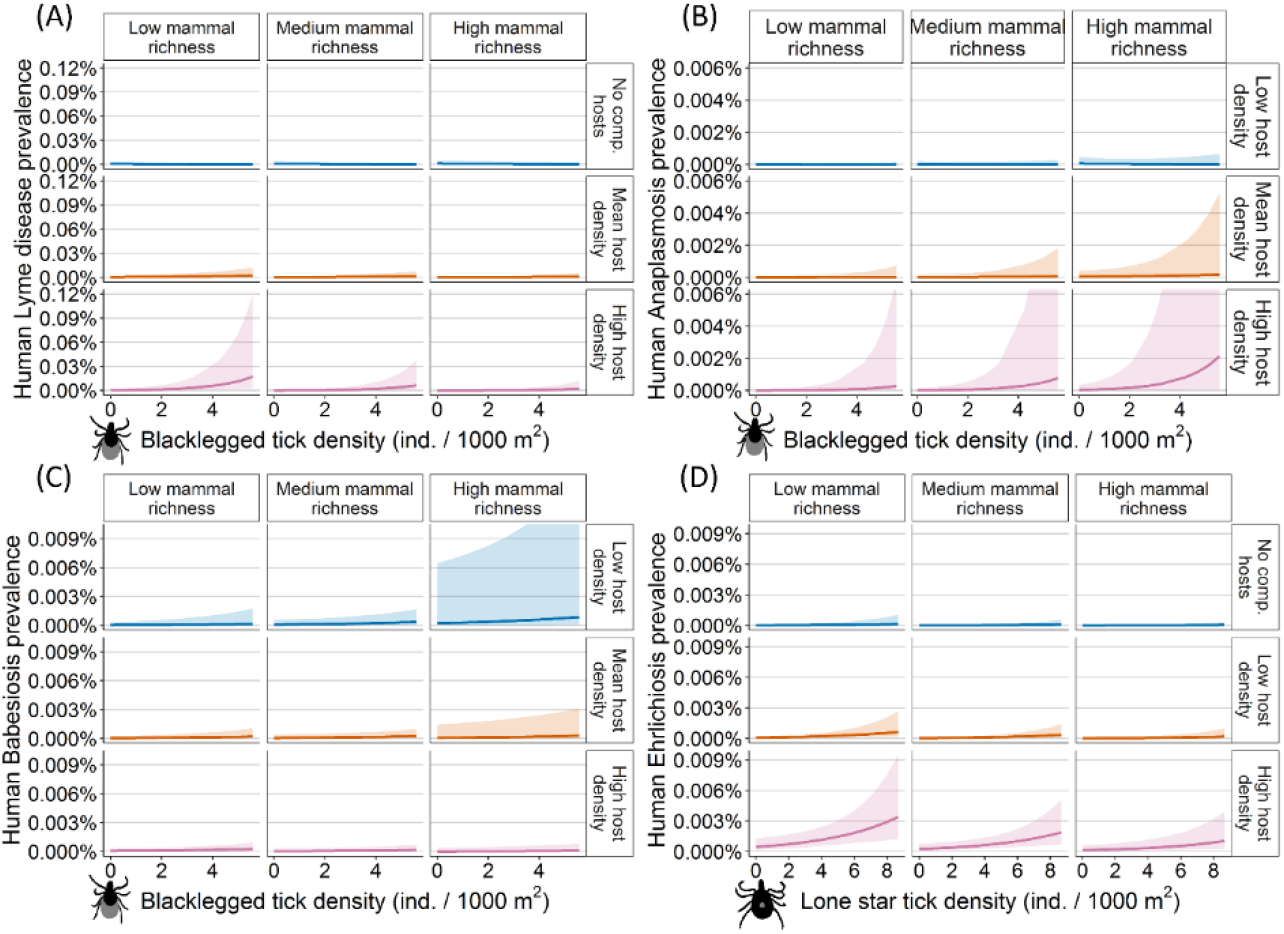
Human prevalence of tick-borne diseases is correlated with reservoir and vector metrics. Model predicted relationships among vector tick density, small mammal community metrics, and human prevalence of disease for A) Lyme disease, B) anaplasmosis, C) babesiosis, and D) ehrlichiosis. For (A) Lyme disease, (B) anaplasmosis, and (C) babesiosis, blacklegged ticks are both *Ixodes scapularis* and *Ixodes pacificus*. Facets of increasing small mammal richness along the top are 5, 8, and 11 species, respectively. For (A) Lyme disease, facets of increasing host density down the right side are 0, 2, and 4 mice per 10000 m^2^, respectively. For (B) anaplasmosis and (C) babesiosis, facets of increasing host density down the right side are 3, 5, and 7 small mammal individuals per 10000 m^2^, respectively. For (D) ehrlichiosis, the facets of increasing host density down the right side are no deer, low deer density, and high deer density. Coloration for reservoir host abundance is consistent across diseases: blue lines are no/low abundance, orange lines are mean abundance, and pink lines are high abundance. Models indicate positive correlations of vector tick density with human prevalence of disease, such that human disease prevalence is predicted to be highest in areas with high vector tick density. For Lyme disease (A), anaplasmosis (B), and ehrlichiosis (D) there was a positive relationship between host abundance and disease prevalence. Accounting for differences in tick and reservoir host abundance, we found significant negative correlations between diversity and disease incidence for Lyme disease (A) and ehrlichiosis (D), but a significant positive correlation between diversity and disease for anaplasmosis (B). The observed patterns are consistent with the dilution effect hypothesis, which posits that a diluting (negative) relationship between diversity and disease is expected when hosts differ in their ability to maintain and transmit pathogens (e.g. Lyme disease and ehrlichiosis); when this condition is not met (e.g. anaplasmosis), an amplifying (positive) relationship is expected.

Because of the opposing direction of the diversity-disease relationship for Lyme disease and anaplasmosis, a non-monotonic relationship emerged between small mammal richness and total tick-borne disease cases, such that human disease incidence was higher at the extremes of small mammal richness (<8 and >14 species per 10,000 m^2^) than at intermediate numbers of species (9–13 species; Fig. 3). A similar pattern emerged when babesiosis and ehrlichiosis were included in the calculation of total human disease, likely due to low incidence of these diseases (Fig. S3). Thus, while mean mammal richness at NEON sites was 8 [7.6, 8.5] species per 10,000 m^2^, our model suggests that maintaining 9-13 species per 10,000 m^2^ should result in the lowest incidence of human tick-borne diseases regardless of tick and host densities (Fig. 3). Specifically, the conservation of small mammal richness to 9-13 species per 10,000 m^2^ is predicted to prevent ~8,600 additional total U.S. tick-borne disease cases annually and an average of 1,000, and as high as 5,700 (when tick and reservoir host densities are high), additional U.S. Lyme disease-caused DALYs annually, relative to 2017 data (Fig. S2).

**Figure 3.**
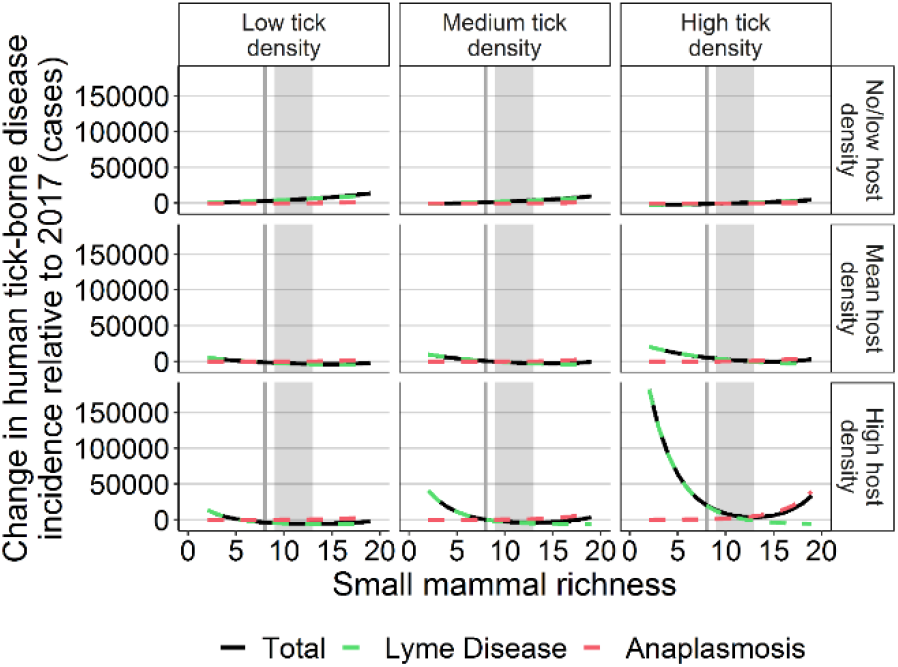
Changes in human tick-borne disease incidence. Changes in human disease incidence for Lyme disease (green lines), anaplasmosis (pink lines), and the sum of the two diseases (total; black solid lines) relative to incidence in the U.S. in 2017. Facets of low, medium, and high tick density along the top are 0.75, 2, and 4 blacklegged ticks (*Ixodes scapularis* and *Ixodes pacificus*) per 1000 m^2^, respectively. Facets of host density down the right are 0, 2, and 4 mice per 10000 m^2^ and 3, 5, and 7 small mammals per 10000 m^2^ for Lyme disease and anaplasmosis respectively. Vertical dark grey line indicated median small mammal richness across NEON sites. Light grey rectangle from 9 to 13 small mammal species represents the mammal richness required to maintain the lowest disease incidence across tick and reservoir host densities. Figures suggest that the magnitude of the relationship between small mammal richness and total tick-borne disease incidence is dependent on the densities of ticks and reservoir hosts. Similarly, figures suggest that the relationship between small mammal richness and total tick-borne disease incidence is non-monotonic and is driven by the reductions in Lyme disease as small mammal richness increases from low (<8 species) to median (8 species), then by the increase in anaplasmosis as small mammal richness increases from median (8 species) to high (>13 species).

Finally, to evaluate the contribution of changes to density of infected ticks on human disease prevalence (Fig. 1), we conducted sequential regressions. For Lyme disease, anaplasmosis, and babesiosis, we found that human disease was positively associated with density of infected ticks (Fig. 4A,D,H), which was driven by the proportion of infected ticks (Fig. 4B,E,I, Table S4). For Lyme disease, the diluting relationship between small mammal diversity and human disease (Fig. 4, Table S4) seemed to be mediated by the effect of small mammal richness on the association between the proportion of infected ticks and reservoir host density (Fig. 4C). When small mammal richness was low, the proportion of infected ticks was positively related to reservoir host density, but this relationship was not different from zero when small mammal richness was high (Fig. 4C). Conversely, the amplifying relationship between small mammal diversity and anaplasmosis and babesiosis was a function of reservoir host density increasing with small mammal richness (Fig. 4G,K), which in turn fueled an increase in the proportion of infected ticks (Fig. 4F,J; Table S4). Despite observing a small mammal dilution effect and a positive effect of deer densities for ehrlichiosis in the model selection analyses that included all sites, we did not find evidence for these effects in the sequential regressions (Table S4), likely because subsetting the data resulted in both reduced samples sizes and the exclusion of sites with no deer.

**Figure 4.**
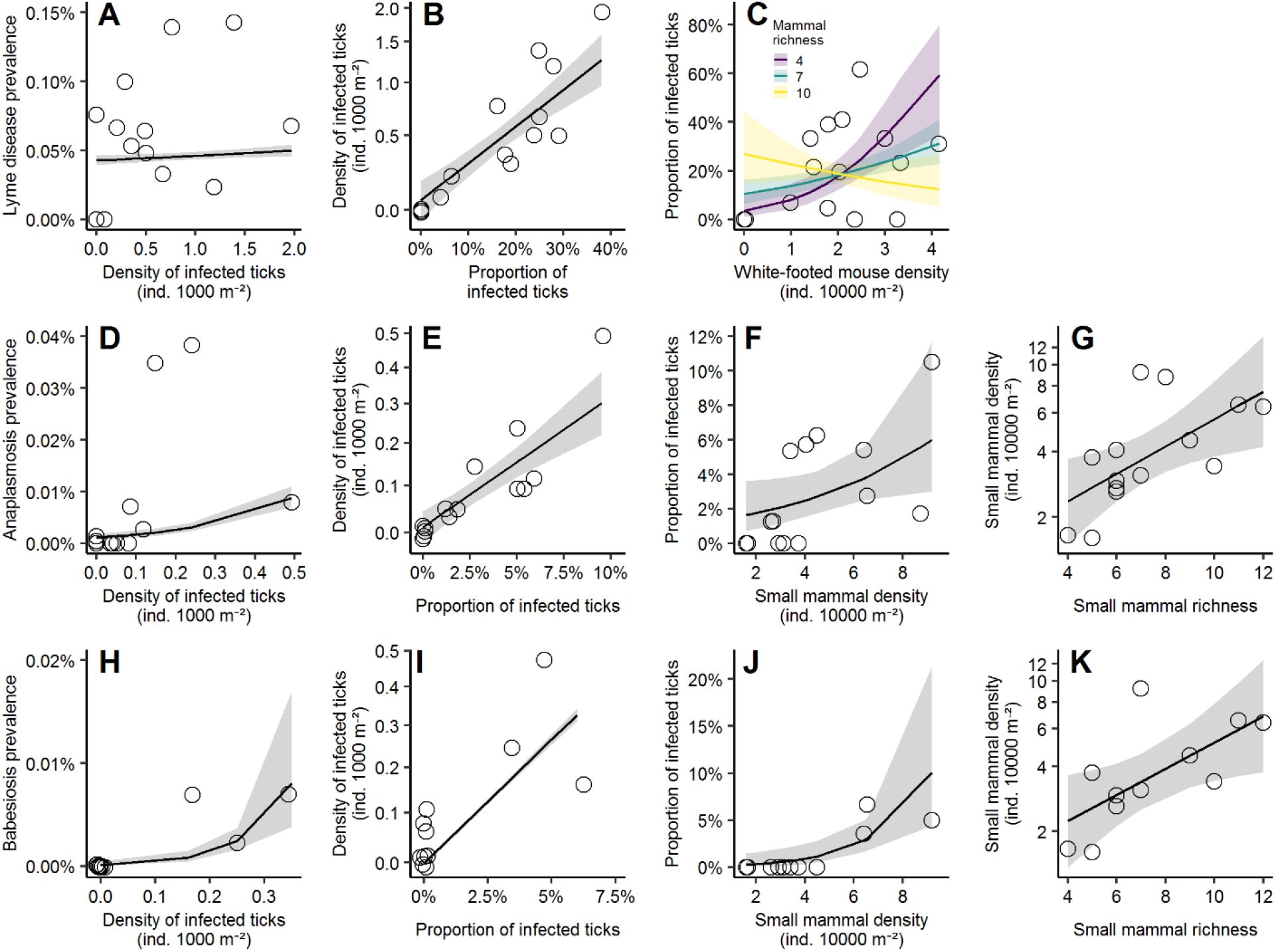
Mammal richness is indirectly correlated with human prevalence of tick-borne diseases. Relationships among (A) human prevalence of Lyme disease and density of *Borrelia burgdorferi*-infected blacklegged ticks; (B) density of infected ticks and proportion of *B. burgdorferi*-infected ticks; (C) proportion of infected ticks, white-footed mouse density, and small mammal richness; (D) human prevalence of anaplasmosis and density of *Anaplasma phagocytophilum*-infected blacklegged ticks; (E) density of infected ticks and proportion of *A. phagocytophilum*-infected ticks; (F) proportion of infected ticks and small mammal density; (G) small mammal density and small mammal richness; (H) human prevalence of babesiosis and density of *Babesia microti*-infected blacklegged ticks; (I) density of infected ticks and proportion of *Ba. microti*-infected ticks; (J) proportion of infected ticks and small mammal density; and (K) small mammal density and small mammal richness. Figures indicate indirect effects of small mammal species richness on human prevalence mediated through increases in proportion of infected ticks, and, in turn, density of infected ticks. Figures support predictions of a negative diversity-disease relationship (dilution) for Lyme disease (A-C), but positive diversity-disease relationships (amplification) for anaplasmosis (D-G) and babesiosis (H-K). Lines for all panels are generalized linear regression coefficients from sequential regressions (see Table S4); ribbons are model indicated standard error. Regression coefficients and statistics are described in Table S. Note: y-axis of (B), (E), (G), (I), and (K) are natural-log transformed. Points on panels are slightly jittered, but do not alter interpretation.

## Discussion

The incidences of tick-borne diseases are increasing globally (1). Because reactive antibiotics can be ineffective at preventing long-term symptoms of these zoonoses if administered too late following infection (7), prevention of these diseases is imperative. Yet, preventative interventions are largely unsuccessful in reducing human disease (1, 8, 9). Here, we related incidence of human tick-borne diseases to tick densities and the densities and diversities of mammalian reservoir hosts across the contiguous U.S. Our results indicate that reservoir host and tick densities are correlated with human disease prevalence for all four diseases. We also found support for our hypothesis that reservoir host richness is associated with human disease prevalence in directions that are predictable by reservoir competence of the host community.

We show that reservoir host and tick densities are correlated with human disease prevalence for all four diseases, likely because as tick and mammal densities increase, so do the density of infected ticks and transmission, as suggested by our sequential regressions. These findings are consistent with previous studies showing positive relationships between reservoir host densities and densities of infected ticks (14, 15). Alternatively, spatial genetic variation in pathogens and host competence could have altered the strength of the relationships among ecological variables and human disease incidence (20, 27). Yet, the inclusion of spatially explicit covariates likely account for much of this potential variation.

Our results indicate that the commonly employed host- and vector-density-targeted interventions should effectively reduce human disease (8, 9); however, these control measures have not translated to fewer cases of human tick-borne diseases in practice (21, 28). The disparity between the expectations from statistical/mathematical models and actual reductions in human disease (21, 28) could be a function of the ecological contexts in which the interventions have been applied, which may have influence over the effectiveness of these interventions, as suggested by our results. For example, in ecological contexts with high Lyme disease risk (low mammal richness, high tick and host densities), our results indicate that the targeted control of both ticks and host densities would synergistically reduce Lyme disease incidence. Conversely, in ecological contexts with moderate Lyme disease risk (median mammal richness, tick density, and host density), our results indicate that the reduction of tick density alone would be most effective in reducing Lyme disease incidence. Therefore, disease interventions targeting ticks would be most effective in reducing Lyme disease, regardless of ecological context, but in high risk contexts, both tick- and host-targeted interventions would prevent the greatest disease incidence. Alternatively, these interventions may be ineffective, because, generally, these models focus on changes to the density of infected ticks (10, 11, 19), which does not directly translate to human disease, because human-tick encounters are the product of both density of infected ticks and human behavior (e.g. repellents, tick checks, etc.) (9, 29).

Because current interventions do not change the species richness of host species, which may be a driver of human disease, an alternative or complementary approach to reducing ticks and reservoir hosts is to conserve or increase the number of reservoir host species. However, this has been a controversial approach to managing tick-borne diseases for several reasons (10, 19, 22, 23, 30), such as concerns that it might increase certain tick-borne diseases while decrease others (10, 19, 22). As predicted, for pathogens with variably competent hosts (i.e. coefficients of variation for reservoirs > 1, Table S3; Lyme disease and ehrlichiosis), human disease prevalence was negatively correlated with small mammal richness, supporting a dilution effect. Specifically, for Lyme disease, the negative relationship between small mammal richness and human disease prevalence increased in magnitude as reservoir host density increased. Also, in agreement with our predictions, human disease prevalence was positively correlated with small mammal richness for anaplasmosis (a pathogen with similarly poor hosts, Table S3), supporting an amplification effect. For babesiosis, the weak positive relationship between small mammal richness and human disease prevalence became negative as reservoir host density increased, supporting an amplification effect. These results imply that determining reservoir competence of a range of host species for vector-borne pathogens may provide valuable insights into whether or not biodiversity conservation would dilute or amplify disease risk (10, 23, 30). Thus, caution must be taken when conserving mammal species as a tool for tick-borne disease control, given the non-monotonic relationship between mammalian richness and total tick-borne disease cases (Fig. 3).

The goal of our study was to evaluate the role of ticks and reservoir host communities in driving broad-scale spatial patterns of human tick-borne diseases, which was not possible until the establishment of NEON. Yet, the ecological data collected by NEON are not without limitations. Specifically, because NEON only employed one type of small mammal sampling (e.g., nocturnal sampling with traps set on the ground), we may be lacking good information on the presence and abundance of some competent reservoir hosts species, resulting in underestimates of host richness and abundance. In using host abundance of the most competent reservoir host for models of Lyme disease and ehrlichiosis, we may be lacking a measure of the abundance of other potentially competent reservoir hosts. Yet, of the species with relatively high realized reservoir competence for *B. burgdorferi* (>0.5; *Peromyscus leucopus* and *Tamias striatus*) (15), *P. leucopus* was found in higher abundances and at more sites, so we chose this species as the main reservoir host for the Lyme disease models. Further, in the western U.S., as the most competent reservoir host (*Sciurus griseus*) (31) was not sampled at NEON sites, we used moderately competent reservoirs (*Peromyscus boylii, P. truei,* and *P. maniculatus*) (31), which may not fully represent the influence of *S. griseus* on *B. burgdorferi* transmission at these sites. Additionally, we recognize that lizards play an important role in regulating transmission of *B. burgdorferi* in the western U.S. (31), but information on the densities of these species at NEON sites was not available, so we are unable to elucidate their role in our models. Finally, in the limited data on reservoir competence for *E. chaffeensis*, the main reservoir hosts are Ruminants (the only native ruminant in the eastern U.S. is white-tailed deer) and Leporidae (rabbits/hares) (17). Yet, because of the high encounter rate between white-tailed deer and *A. americanum* (i.e. reservoir potential) (17), white-tailed deer are still likely the major reservoir hosts for *E. chaffeensis*, and as such, we selected this species as the reservoir host for the ehrlichiosis models.

Additionally, the tick sampling methodology used by NEON may limit detections of nymphal *I. scapularis* in the southeastern U.S. (32); thus, despite the regular sampling of nymphal ticks at these sites, tick densities at these sites might be underestimated. Importantly, although nymphal and adult ticks play different roles in human disease, the use of nymphal ticks instead of pooling all tick stages does not appreciably alter the results nor interpretation of our analyses (Table S5). Similarly, including NEON sites without forest land-cover in our analyses, and, thus without the ecotonal habitat of the focal tick species, could have generated spurious patterns, but removing these sites from the analyses did not change the results nor interpretation of our analyses (Table S6). Thus, we believe that the established patterns in our results are robust and ecologically sound.

As tick-borne disease incidence continues to rise across the U.S. (2, 4), preventative measures, such as controlling ticks and reservoir host densities and diversity, are essential to reduce human tick-borne disease. While individual relationships among reservoir hosts, ticks, density of infected ticks, and human disease incidence have been previously supported (11, 15, 19, 20, 22), our study is the first to support all links among these variables at broad spatial scales, supporting the promise of proactive rather than reactive approaches to tick-borne disease management. Although host- and vector-targeted interventions reduce densities of infected ticks (33–35), the individual use of these interventions to reduce human disease have shown some shortcomings in practice (21, 28, 33, 35, 36). Our results indicate that the ecological context (tick and host densities and host diversity) may influence the relative effectiveness of these control measures. Further, an alternative and/or complimentary approach to traditional tick-borne disease control may be the conservation of small mammal species within a feasible but specific range of richness levels. Consequently, future work should investigate the effectiveness of control measures targeting tick and reservoir host densities across a mammalian host richness gradient to determine what levels and combinations of interventions would be most effective at preventing human tick-borne diseases.

## Materials and Methods

### Study area

Our analyses paired site-level estimates of tick density, tick-borne pathogen prevalence, and small mammal communities from 38 climatically and ecologically variable sites in the National Ecological Observatory Network (NEON; Figure S1; Table S2) with county-level human case counts of tick-borne diseases collected by the Center for Disease Control (CDC) Notifiable Diseases Surveillance System (NNDSS) and Division of Parasitic Diseases and Malaria (DPDM). The study area included 35 counties in 21 states across the contiguous United States (Table S2). Due to variability in data collection across NEON sites and availability of disease incidence data, not all sites or counties were included in each year of the data (see Table S2 for site-year replicates).

### NEON Data

Sampling protocols for ticks, tick-borne pathogens, and small mammals in this section will be described briefly, as detailed information on sampling protocols are available through the appropriately cited NEON sampling protocols (37, 38). NEON data used in this study were downloaded on 1 November 2019; datasets used in this study are outlined in Table S7.

#### Tick sampling

Starting in 2014, at each site, tick sampling occurred in six plots that were iteratively sampled. Sampling frequency was dependent on whether ticks were detected. Sampling began with a sampling events every six weeks, but collection of one or more ticks prompted sampling events every three weeks (37). The start of sampling at a site coincided within two weeks of the onset of vegetation green-up and ended within two weeks of senescence (typically April-September, but may be March-October depending on site and weather). Each tick sampling plot was 40 x 40m; the perimeter of each sampling plot was sampled with a 1 x 1m drag cloth. If vegetation within a sampling plot was too thick to allow dragging, flagging was used either instead of dragging or in conjunction with dragging. Ticks were identified to species and life stage. For each focal species and sampling event, nymph and adult abundances were pooled. Tick abundances were converted to densities based on area sampled (individuals per m^2^) and scaled to individuals per 1,000 m^2^. We then calculated the density of focal tick species (*Ixodes scapularis*, *Ixodes pacificus*, or *Amblyomma americanum*) as the mean number of individuals collected per sampling event per sampling plot, to account for differences in the number of tick sampling events across sites and years. See Fig. S1 for NEON sites at which each focal tick species was sampled. Ticks were not supplementally sampled from small mammal hosts.

#### Tick-borne pathogen prevalence

Testing of pathogen prevalence in ticks occurred at 13 sites in the eastern U.S. (Table S8), starting in 2014 (37). At the time of data analysis, tick pathogen prevalence was only available for 2014-2017. At a given site, a subset of sampled nymphal ticks were tested annually for the presence of zoonotic pathogens (Table S8). *Ixodes scapularis* (eastern blacklegged ticks) were tested for *Anaplasma phagocytophilum*, *Babesia microti*, and *Borrelia burgdorferi*. *Amblyomma americanum* (lone star ticks) were tested for *Ehrlichia chaffeensis*. Pathogens status in nymphal ticks was tested using next-generation sequencing and 16S rRNA primers. As quality control, we excluded all pathogen status results that did not also test positive for hard-tick DNA. Pathogen prevalence at a site was estimated as the proportion of nymphal ticks that tested positive for a given pathogen.

#### Small mammal trapping

Starting in 2014, at each site, trapping plots were arranged in three to eight plots of 100 live traps (Sherman) arranged in a 10 x 10 grid, with 10 m spacing (100 x 100m area) (38). Trapping plots were separated by at least 135 m. NEON field technicians trapped, identified, and released small mammals from each grid either one or three nights (depending on whether sampling for diversity or pathogens, respectively) per month or every other month (depending on site designation as core or relocatable, respectively) during the growing season within a 21 day window centered on the new moon (typically April-September, but may be March-October depending on site and weather). Ethical approval was obtained from IACUC (38).

Small mammal richness was the total number of unique species collected across all sampling plots at a given site each year. Small mammal abundance for a given site each year was calculated as the mean number of individuals of all species collected per trap night per sampling plot (individuals per 10,000 m^2^), to account for differences in the number of sampling events and plots across sites and years. Because captured individuals were marked, we excluded recaptures from the estimates of small mammal abundance. See Table S9 for species by site matrix. Similarly, the abundance of main reservoir species for *B. burgdorferi,* the white-footed mouse (*Peromyscus leucopus*) in the eastern US and the brush mouse (*Peromyscus boylii*), the pinyon mouse (*Peromyscus truei*), and the deer mouse (*Peromyscus maniculatus*) at sites located within the range of *Ixodes pacificus* (western blacklegged tick) in the western U.S. (NEON sites: ABBY, ONAQ, SOAP, and SJER), was calculated as the mean number of individuals collected per trap night per sampling plot (individuals per 10,000 m^2^). The abundances of these four species were pooled and termed “*B. burgdorferi* reservoir density”.

### Deer density estimates

As white-tailed deer are the assumed main reservoir host for *E. chaffeensis* (17, 39), we included deer densities as a predictor of ehrlichiosis. The most recent white-tailed deer density estimates for the United States cover 2001-2005, were compiled by the Quality Deer Management Association, and are hosted by U.S. Forest Service (40). Deer densities were not meant to be a measure of absolute white-tailed deer density, but rather were meant to provide an estimate for relative densities across the continental United States. Therefore, we grouped deer density estimates into three categories: No Deer, Low Density, and High Density. The “No Deer” category represents areas where white-tailed deer are absent, the “Low Density” category represents deer densities (<11.6 deer/km^2^) that are below or on the cusp of ecologically damaging, and the “High Density” category represents deer densities (>11.6 deer/km^2^) that are ecologically damaging and are greatly above historic, pre-European settlement densities (41).

### Human cases of tick-borne diseases

We obtained the annual number of reported cases of Lyme disease, anaplasmosis, ehrlichiosis caused by *E. chaffeensis*, and babesiosis at the county level from the CDC NNDSS and DPDM from 2014 – 2018. At the time of analysis, Lyme disease and babesiosis cases were only available from 2014-2017. Tick-borne disease case definitions by the CDC includes both confirmed and probable cases, to address under-reporting (42, 43). Due to differences in Lyme disease case definitions between the CDC and Massachusetts Department of Health starting in 2016, most cases for Massachusetts are not reported to the CDC (42, 44); thus, case data from Worcester, Massachusetts was limited to 2014 and 2015. Anaplasmosis, ehrlichiosis, and babesiosis are not reportable conditions for all states in all years: anaplasmosis and ehrlichiosis was not reported in Colorado and New Mexico across years, while babesiosis was not reported in Arizona, Colorado, Georgia, Kansas, New Mexico, and Oklahoma across years. Further, babesiosis was not reported from Virginia and Florida before 2017, so data from sites in these states before 2017 were not included in the analyses. Tick-borne disease cases are reported in the patient’s county of residence, so reporting errors due to travel are possible, especially in non-endemic and non-emerging areas.

### Statistical analyses

All statistical analyses were conducted in R version 3.6.1 (45). For the national-scale analyses, we used model selection and generalized linear mixed effects models (GLMMs) with a binomial distribution to explore patterns among human cases of tick-borne disease prevalence, tick densities, small mammal diversity, and reservoir species abundance. Responses for each model were a binary county-level tick-borne disease case count and population. For Lyme disease and anaplasmosis models, we used glmer in the lme4 package (24) and for the ehrlichiosis and babesiosis models, we used bglmer in the blme package (46) with a gamma covariance prior, due to issues of singular fit related to low incidence of these diseases. Predictor variables included tick densities, reservoir host abundance, and small mammal richness. For Lyme disease, anaplasmosis, and babesiosis, tick densities were the densities of the eastern (*I. scapularis*) and western (*I. pacificus*) blacklegged ticks. For ehrlichiosis, tick densities were the densities of the lone star tick (*A. americanum*). For the measure of reservoir host abundance, we used the abundance of the primary competent reservoir host. Specifically, for Lyme disease we used the abundance of the eastern reservoir (white-footed mice) and the western mammalian reservoirs (pinyon, brush, and deer mice). For ehrlichiosis, we used the abundance of white-tailed deer for the measure of reservoir host abundance. For both anaplasmosis and babesiosis (diseases with evenly poor reservoir hosts, Table S1 (15, 18)), we used abundance of all small mammals for the measure of host reservoir abundance. County was included as a random term in all models, because annual observations within the same county are not independent.

We conducted model selection in which we fit all possible combinations of main effects and biologically relevant two-way interactions of all biotic variables (interactions between host density and tick density and between host density and host richness; dredge function in MuMIn R package) (47). To account for differences in questing height of blacklegged ticks along a north-south gradient (25), we included latitude as a covariate for blacklegged tick-borne diseases. Further, to account for potential climate-related differences in relationships between wildlife variables and human disease incidence (20), all models included covariates of mean annual temperature and annual precipitation. Analysis of correlations among of variables indicated correlation >0.6 for only a single pair of variables that jointly appear in any GLMMs: temperature and latitude (Fig. S4). Despite this correlation, the inclusion of both variables in models is important to capture known ecological/behavioral gradients along each variable. To find best fitting models, we used a combination of the lowest Akaike’s Information Criterion with bias-correction (AICc) and the law of parsimony; such that competing models (ΔAICc < 2) with fewer degrees of freedom than models with lowest AICc were selected as the best model. Likelihood ratio tests against a null model containing only the random term of county and fixed effects of mean annual temperature and annual precipitation (and latitude for blacklegged tick-borne disease models) were used to determine overall p-values for best models. We calculated AICc weights (*w*) and marginal (fixed) and conditional (fixed and random effects) R^2^ values (48). We performed model diagnostics on residuals of best models using the DHARMa R package (49), which indicated model assumptions had been met and no spatial autocorrelation.

For models with significant effects of small mammal richness on human disease prevalence, we calculated the differences in case prevalence from 2017 (the year with the most recent complete data) by subtracting the median state-level prevalence for a given disease in 2017 from best model-predicted prevalence. We then multiplied this relative change in prevalence by the U.S. population from all states reporting anaplasmosis cases to get total change in disease incidence across the U.S. These are reported in Figure 3 in the main text. Change in Lyme disease incidence was then converted to change in annual disability-adjusted life years (DALYs) to provide an estimate of overall disease burden from changing disease incidence. Average case DALYs for Lyme disease have been estimated for patients with different Lyme disease outcomes: erythema migrans (0.005 DALYs), disseminated Lyme disease (0.113 DALYs), and Lyme-related persisting symptoms (1.661 DALYs) (26). Relative prevalence of these outcomes per Lyme disease diagnosis are 82.8% for erythema migrans, 8.6% for disseminated LD, and 8.6% for persisting Lyme disease symptoms (7). Thus, the estimated DALYs per Lyme disease case is 0.156.

To address our second objective of direct and indirect effects of tick density and reservoir host metrics on human disease prevalence, we used sequential regressions rather than traditional structural equation models (SEMs), because of limited data on tick infection prevalence (<30 replicates per pathogen). Models were fit using generalized linear models with either a normal error distribution (for tick and mammal density models) or a binomial error distribution (for tick and human disease prevalence models). All models were hypothesized *a priori* from national-scale analyses, the ecology of these pathogens, and previous studies (10, 19, 23, 50); see Fig. 1 in main text for *a priori* hypothesized links. Significance of relationships was found with type 3 error and p-values were adjusted using the Holm-Bonferroni sequential correction (51), by which *k* relationships are ranked (*i*) by their p-values (*Pi*) and p-values are adjust by the equation: (*k* − *i* + 1) * *P*_*i*_. Significance levels of all models was *P* < 0.05.

## Acknowledgements

We would like to thank the National Ecological Observatory Network for collection and maintenance of ecological data. We would like to thank K.N.H. and S.P.H. at the Centers for Disease Control for providing data access. We thank the Rohr lab and S.L. Rumschlag for feedback on this manuscript, and A.M. Kilpatrick for discussions on community competence and tick-borne diseases. Funding was provided by the University of Notre Dame and the National Science Foundation (DEB-1518681, EF-1241889, IOS-1754868).

