## Supplement for "Reservoir host community and vector density predict human tick-borne diseases across the United States"

**Supplemental Information; Text, Tables and Figures**

### Supplemental Text

Diversity-Disease Relationships

Zoonotic disease transmission to humans entails complex interactions between vectors and hosts (1–4) that are influenced by a variety of climatic factors (e.g. humidity, extreme temperatures, and snowpack depth) (5–7) and host community characteristics (e.g. relative abundance, identity, diversity, and reservoir competency of community members) (8–12). In particular, host community diversity can strongly influence pathogen transmission dynamics from hosts to vectors and vectors to humans (4, 8, 13, 14). Mechanistic understanding of these diversity-disease relationships assumes 1) hosts differ in their ability to maintain and transmit pathogens (reservoir competency), because parasites are likely to adapt to infect abundant species and abundant hosts invest in reproduction and growth at the expense of parasite defense, 2) abundant hosts are more likely to be persistent through time in an area, and 3) as species are introduced into an ecosystem, this reduces the abundance of common, competent hosts (4, 14, 15). When these conditions are met, a diluting effect of diversity on disease (negative relationship between diversity and disease) is expected (4, 14). Yet, when these conditions are not met, an amplifying effect of diversity (positive relationship between diversity and disease) or no detectable effect of diversity on disease is likely (4, 13, 16, 17). Thus, the reservoir competence of hosts can influence the direction of diversity-disease relationships (see Fig. 1 in main text).

### Supplemental Text References

### Attribution of Silhouette Images

Silhouettes of organisms used throughout the manuscript are presented in accordance with licensing agreements. Below, we provide information on the creators' contributions to the images. Licensing agreements include: Public Domain (https://creativecommons.org/publicdomain/mark/1.0/), Creative Commons Attribution 3.0 Unported license (https://creativecommons.org/licenses/by/3.0/), Attribution-ShareAlike 4.0 International (https://creativecommons.org/licenses/by-sa/4.0/), and Attribution-ShareAlike 4.0 International (https://creativecommons.org/licenses/by-sa/4.0/).

*Tamias striatus*: created by MB Mahon (vectorization), R Hodnett (photography), CC BY-SA 4.0
*Peromyscus leucopus*: created by MB Mahon (vectorization), DGE Robertson (photography), CC BY-SA 3.0
*Blarina brevicauda*: created by MB Mahon (vectorization), Fralambert (photography), CC BY-SA 4.0
*Sciurus carolinensis*: created by MB Mahon (vectorization), T Friedel (photography), CC BY 3.0
*Sorex cinereus*: created by MB Mahon (vectorization), J Bachman (print), Public Domain
Woman Silhouette: created by MB Mahon (adjustments), Openclipart (vectorization), Public Domain
Man Silhouette: created by MB Mahon (adjustments), RexxS (vectorization), Public Domain
*Amblyomma americanum*: created by MB Mahon (vectorization), J Gathany (photography), Public Domain
*Ixodes scapularis*: created by MB Mahon (vectorization), J Gathany (photography), Public Domain

**Supplemental Tables and Figures**

Table S1. Mean reservoir competences for the four focal tick-borne pathogens in this study. Reservoir competence values come from refs. (2, 18, 19). The coefficient of variation (CV) is calculated as the standard deviation/mean of the reservoir competences of all tested reservoir hosts, and, thus, is the relativized standard deviation accounting for differences in the mean. Pathogens with CV values > 1 are considered those with "variably competent hosts", while pathogens with CV values <1 are considered those with "similarly competent hosts". Our approach of estimating the mean and variance of reservoir competence is limited by the number of species that have had their reservoir competence estimated, which is only a subset of the small mammal species that are likely to be exposed to each of these pathogens.

| Pathogen | Mean Reservoir Competence† | C.V. | Data source |
|---|---|---|---|
| *Borrelia burgdorferi* | 0.244 | 1.28 | Ostfeld et al. 2018 |
| *Ehrlichia chaffeensis* | 0.018 | 1.099 | Allan et al. 2010 |
| *Babesia microti* | 0.137 | 0.811 | Ostfeld et al. 2018 |
| *Anaplasma phagocytophilum* | 0.035 | 0.804 | Keesing et al. 2014 |

†Reservoir competences for *B. burgdorferi*, *B. microti*, and *A. phagocytophilum* are realized reservoir competence (proportion of ticks that are infected following a feeding on infected hosts). Reservoir competences for *E. chaffeensis* are estimates of reservoir capacity (absolute contribution of a reservoir host to pathogen infection prevalence in tick populations).

Table S2. List of NEON sites, associated County and State locations, and years of data included in the analyses. For a site-year replicate to be included in the GLMM analyses, all relevant data needed to be available: human disease prevalence, reservoir host density and diversity, and tick density.

| State | County | Years Included | NEON Site |
|---|---|---|---|
| Alabama | Choctaw | 2016-2018 | LENO |
| Alabama | Hale | 2014-2018 | TALL |
| Alabama | Marengo | 2015-2018 | DELA |
| Arizona | Pima | 2016-2018 | SRER |
| California | Fresno | 2018 | SOAP |
| California | Madera | 2016-2018 | SJER |
| Colorado | Boulder | 2016-2018 | NIWO |
| Colorado | Boulder | 2017-2018 | RMNP |
| Colorado | Logan | 2014, 2016-2018 | STER |
| Colorado | Weld | 2014, 2016-2018 | CPER |
| Florida | Polk | 2014-2018 | DSNY |
| Florida | Putnam | 2014-2018 | OSBS |
| Georgia | Baker | 2014-2018 | JERC |
| Kansas | Jefferson | 2015-2018 | UKFS |
| Kansas | Riley | 2017-2018 | KONA |
| Kansas | Riley | 2015-2018 | KONZ |
| Maryland | Anne Arundel | 2015-2018 | SERC |
| Massachusetts | Worcester | 2014-2015 | HARV |
| Michigan | Gogebic | 2014-2018 | UNDE |
| New Hampshire | Carroll | 2014-2018 | BART |
| New Mexico | Dona Ana | 2016-2018 | JORN |
| North Dakota | Morton | 2016-2018 | NOGP |
| North Dakota | Stutsman | 2017-2018 | DCFS |
| North Dakota | Stutsman | 2014-2018 | WOOD |
| Oklahoma | Washita | 2015-2018 | OAES |
| Tennessee | Anderson | 2014-2018 | ORNL |
| Tennessee | Sevier | 2015-2018 | GRSM |
| Texas | Wise | 2016-2018 | CLBJ |
| Utah | San Juan | 2015-2018 | MOAB |
| Utah | Tooele | 2014, 2016-2018 | ONAQ |
| Virginia | Clarke | 2015-2018 | BLAN |
| Virginia | Giles | 2017-2018 | MLBS |
| Virginia | Warren | 2014-2018 | SCBI |
| Washington | Clarke | 2016-2018 | ABBY |
| Washington | Skamania | 2018 | WREF |
| Wisconsin | Lincoln | 2016-2018 | TREE |
| Wisconsin | Price | 2016-2018 | STEI |
| Wyoming | Park | 2018 | YELL |

Figure S1. Target tick species (*Amblyomma americanum*, *Ixodes scapularis*, and *Ixodes pacificus*) found at 38 terrestrial National Ecological Observatory Network (NEON) sites across the continental United States. Red circles are sites at which only *Amblyomma*, *americanum* were found; green circles are sites at which only *Ixodes scapularis* were found; yellow circles are sites at which both *A. americanum* and *I. scapularis* were found; light blue circles are sites at which only *Ixodes pacificus* were found; and "X"s are sites at which no target tick species were found. Note: NEON site locations have been slightly jittered to minimize overlap between spatially close sites.

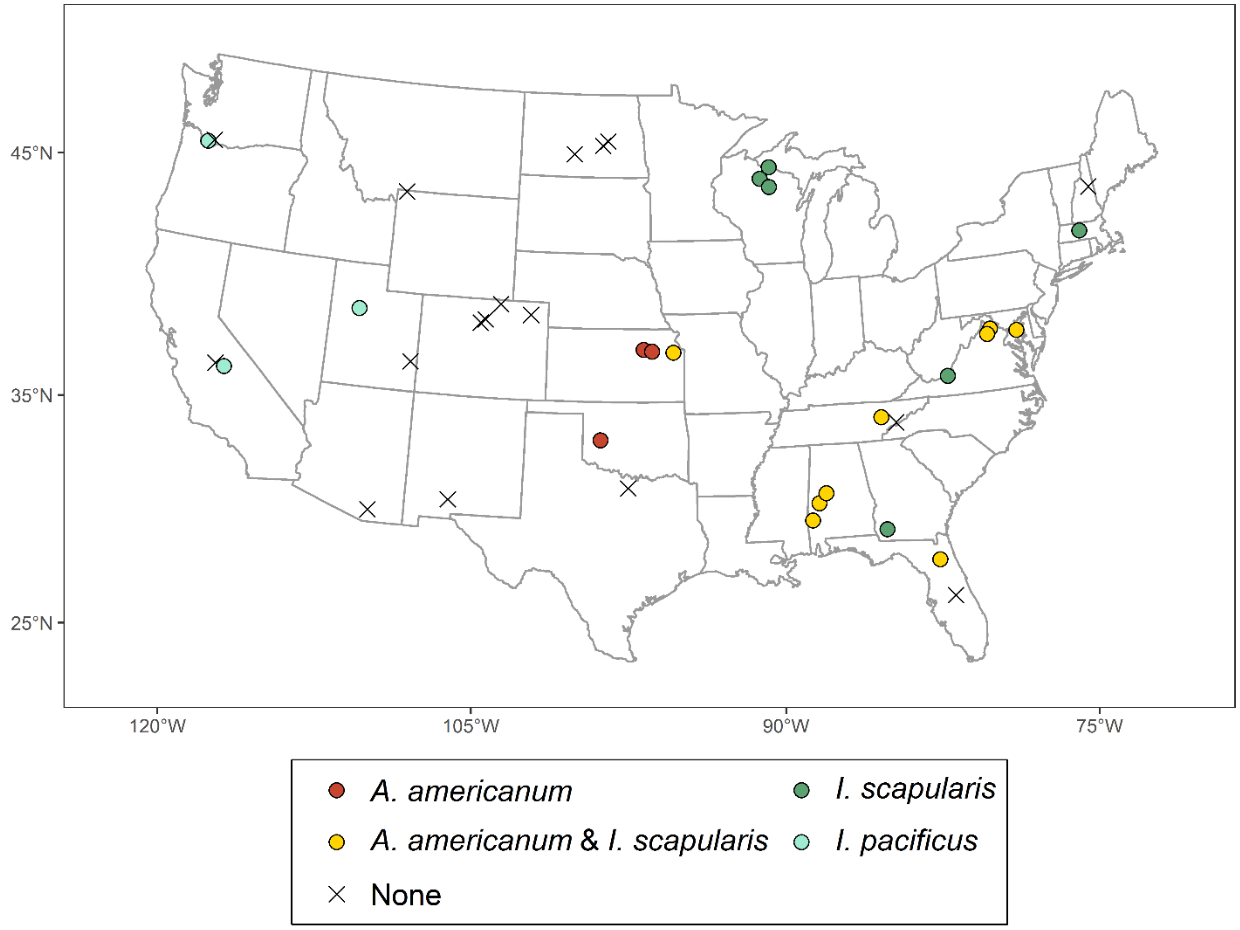

Table S3. Best and competing models (ΔAICc < 2) for each disease. Predictor variables are vector density (eastern and western blacklegged ticks – Lyme Disease, Anaplasmosis, and Babesiosis; lone star ticks – Ehrlichiosis), reservoir density (*B. burgdorferi* reservoir abundance – Lyme Disease; small mammal abundance – Anaplasmosis and Babesiosis; white-tailed deer – Ehrlichiosis), and small mammal richness. Model selection allowed for the main effect of all predictors and two-way interactions between vector and reservoir density and between reservoir density and small mammal richness. All models included a random effect of county. All models included covariates of mean annual temperature and total annual precipitation. Models for blacklegged tick-borne diseases (Lyme Disease, Anaplasmosis, and Babesiosis) also included covariate of latitude. Models for Lyme Disease also included CDC reporting type (definition changed in 2017).

| Model/Variables | df | logLik | AICc | ΔAICc | *w* | Marg/Cond $R^2$ |
|---|---|---|---|---|---|---|
| Lyme Disease | | | | | | |
| **Blacklegged tick density * *B. burgdorferi* reservoir density + Small mammal richness * *B. burgdorferi* reservoir abundance** | **11** | **-183.99** | **393.17** | **0** | **0.99** | **0.18/0.77** |
| Anaplasmosis | | | | | | |
| **Blacklegged tick density * Small mammal density + Small mammal richness** | **9** | **-91.84** | **203.39** | **0** | **0.69** | **0.32/0.82** |
| Babesiosis | | | | | | |
| **Blacklegged tick density + Small mammal density * Small mammal richness** | **9** | **-26.9** | **75.63** | **0** | **0.51** | **0.73/0.74** |
| Ehrlichiosis | | | | | | |
| Lone star tick density * Deer Density + Small mammal richness | 8 | -52.44 | 122.23 | 0 | 0.36 | 0.51/0.53 |
| **Lone star tick density + Deer Density + Small mammal richness** | **7** | **-54.06** | **123.15** | **0.92** | **0.22** | **0.44/0.46** |
| Lone star tick density * Deer Density + Deer Density * Small mammal richness | 9 | -52.00 | 123.70 | 1.47 | 0.17 | 0.49/0.51 |

Figure S2. Changes in human disease burden for Lyme Disease. Change in annual Lyme Disease related disability-adjusted life years (DALYs) relative to DALYs in the U.S. in 2017. Facets of low, medium, and high small mammal richness across the top are 5, 8, and 11 species, respectively. Facets of no competent hosts, mean host density, and high host density across the right side are 0, 2, and 4 mice per 10000 m$^2$, respectively. Coloration of lines follows reservoir host density: blue lines are no/low abundance, orange lines are mean abundance, and pink lines are high abundance. Figure suggests conservation management of mammal species richness is most impactful on human disease burden in areas of high tick and reservoir density.

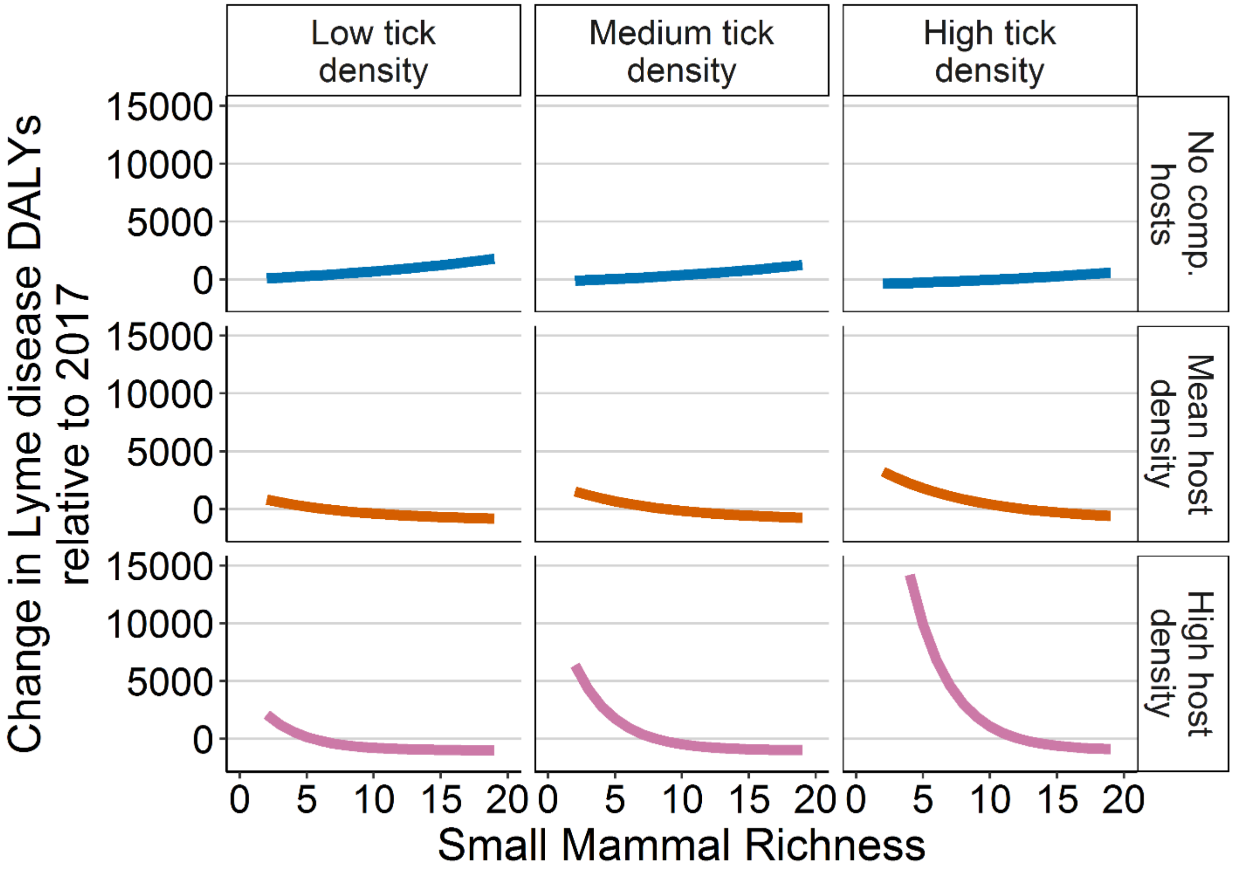

Figure S3. Changes in human disease incidence for four tick-borne diseases. Changes in disease incidence for Lyme Disease (green lines), Anaplasmosis (pink lines), Ehrlichiosis (purple lines), Babesiosis (yellow lines) and the sum of the four diseases (total; black solid lines) relative to incidence in the U.S. in 2017. Facets of tick density along the top are 0.75, 2, and 4 ticks per 1000 $m^2$, respectively. Facets of increasing host density down the right side are 0, 2, and 4 mice per 10000 $m^2$ for Lyme Disease; 3, 5, and 7 small mammal individuals per 10000 $m^2$ for Anaplasmosis and Babesiosis; and no deer, low deer density, and high deer density for Ehrlichiosis. Vertical dark grey line indicated median small mammal richness across NEON sites. Light grey rectangle from 9 to 13 small mammal species represents the mammal richness required to maintain the lowest disease incidence across tick and reservoir host densities. Figures suggest that changes in total human disease cases are driven by changes in Lyme Disease and Anaplasmosis.

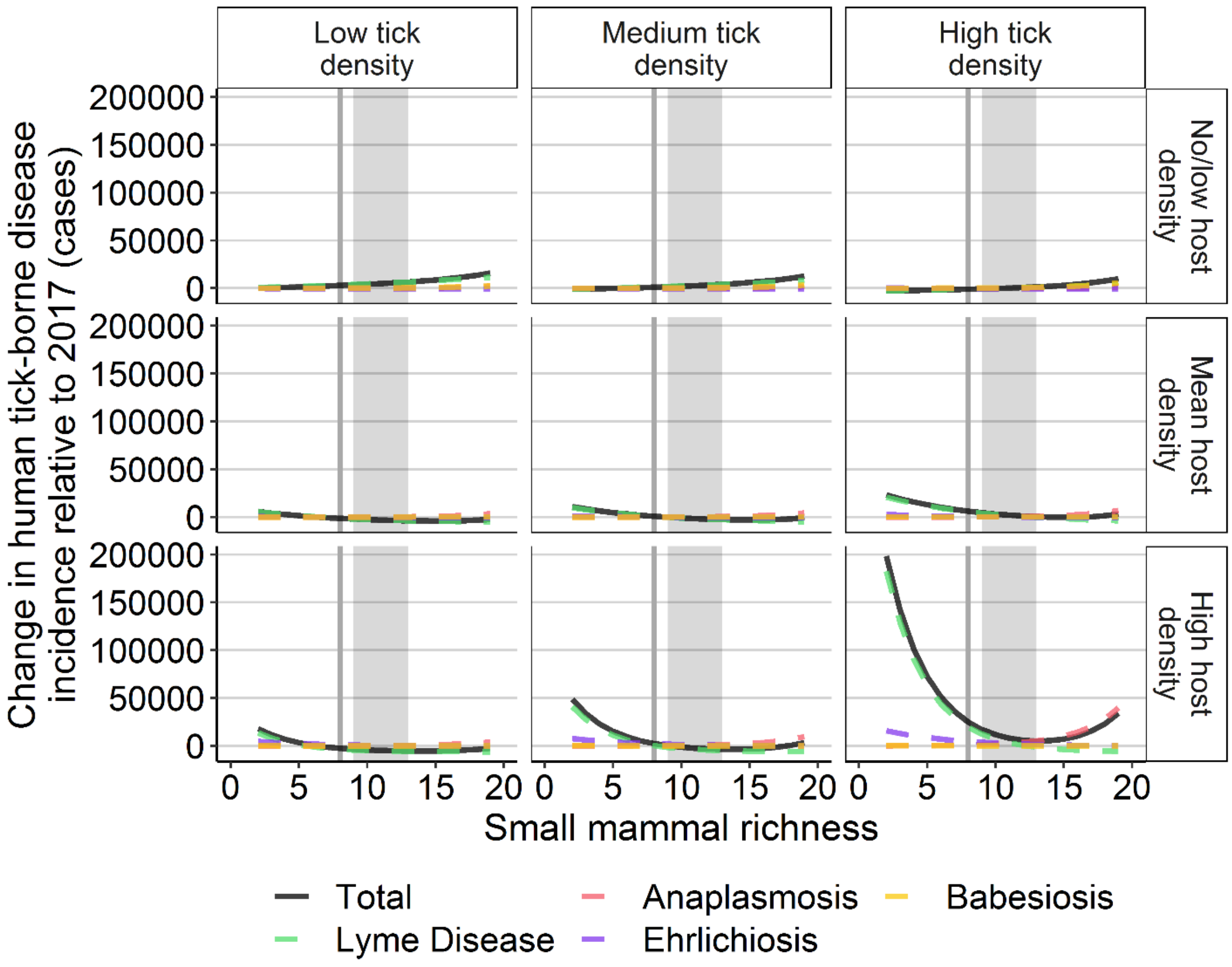

Table S4. Sequential regressions of hypothesized linkages between small mammal richness and human prevalence of tick-borne diseases. Model predictors are on the left, model responses are on the right. Estimates are non-standardized regression coefficients and standard error from generalize linear regressions (binomial distribution for proportion models – human prevalence and tick infection prevalence; Gaussian distribution for all other models). P-values are Holm-Bonferroni sequential corrected p-values (see methods for details). For Lyme Disease, Anaplasmosis, and Babesiosis, we see support for small mammal richness affecting human disease prevalence, mediated through changes to proportion of infected ticks and density of infected ticks. For Ehrlichiosis, we do not see support for this linkage, which may partially be due to not having sites with no deer and minimal knowledge about the ecology of this pathogen.

| Disease / Model | Coefficient Estimate | Test Stat (^F/*$X^2$) | P | R2 |
|---|---|---|---|---|
| Lyme Disease | | | | |
| Density Infected Ticks -> Disease Prop | 0.56 (0.05) | 133.48* | 0.009 | 0.36 |
| Density of Ticks -> Density Infected Ticks | 0.11 (0.02) | 56.70^ | 0.007 | 0.96 |
| Prop. Infected Ticks -> Density Infected Ticks | 1.49 (0.20) | 56.34^ | 0.008 | 0.96 |
| White-footed Mouse Abundance -> Density of Ticks | 0.07 (0.10) | 0.62^ | 0.448 | 0.28 |
| Non-competent Host Abundance -> Density of Ticks | 0.13 (0.06) | 5.65^ | 0.111 | 0.28 |
| White-footed Mouse Abundance -> Prop. Infected Ticks | 1.51 (0.49) | 9.54* | 0.012 | 0.17 |
| Mammal Richness -> Prop. Infected Ticks | 0.35 (0.13) | 6.92* | 0.045 | 0.17 |
| Non-competent Host Abundance -> Prop. Infected Ticks | 0.05 (0.06) | 0.87* | 0.700 | 0.17 |
| Mouse Abundance*Mammal Richness -> Prop. Infected Ticks | -0.17 (0.07) | 6.50* | 0.044 | 0.17 |
| Anaplasmosis | | | | |
| Density Infected Ticks -> Disease Prop | 4.17 (0.56) | 71.71* | 0.006 | 0.37 |
| Density of Ticks -> Density Infected Ticks | 0.03 (0.01) | 12.64^ | 0.016 | 0.88 |
| Prop. Infected Ticks -> Density Infected Ticks | 2.65 (0.40) | 43.66^ | 0.005 | 0.88 |
| Small Mammal Abundance -> Density of Ticks | 0.12 (0.04) | 7.26^ | 0.038 | 0.38 |
| Small Mammal Abundance -> Prop. Infected Ticks | 0.18 (0.09) | 3.92* | 0.048 | 0.09 |
| Mammal Richness -> Small Mammal Abundance | 0.14 (0.05) | 8.01^ | 0.045 | 0.40 |
| Ehrlichiosis | | | | |
| Density Infected Ticks -> Disease Prop | -1.70 (6.07) | 0.080* | 1.000 | 0.01 |
| Density of Ticks -> Density Infected Ticks | 0.04 (0.01) | 9.12^ | 0.020 | 0.78 |
| Prop. Infected Ticks -> Density Infected Ticks | 4.20 (0.53) | 63.63^ | 0.006 | 0.78 |
| Deer Density -> Density of Ticks | -0.21 (0.21) | 1.09^ | 0.924 | 0.04 |
| Deer Density -> Prop. Infected Ticks | -1.68 (0.66) | 8.807* | 0.015 | 0.15 |
| Small Mammal Richness -> Prop. Infected Ticks | 0.01 (0.16) | 0.065* | 0.799 | 0.15 |
| Babesiosis | | | | |
| Density Infected Ticks -> Disease Prop | 11.89 (2.80) | 32.64* | 0.005 | 0.63 |
| Density of Ticks -> Density Infected Ticks | -0.01 (0.01) | 0.41^ | 0.538 | 0.99 |
| Prop. Infected Ticks -> Density Infected Ticks | 4.73 (0.11) | 1954.25^ | 0.006 | 0.99 |
| Small Mammal Abundance -> Density of Ticks | 0.17 (0.04) | 16.14^ | 0.009 | 0.64 |
| Small Mammal Abundance -> Prop. Infected Ticks | 0.48 (0.15) | 11.32* | 0.004 | 0.38 |
| Mammal Richness -> Small Mammal Abundance | 0.14 (0.05) | 7.59^ | 0.044 | 0.46 |

Table S5. Regression coefficients and statistics from best-fit models when nymphal ticks are used instead all ticks. Predictor variables were blacklegged tick density, lone star tick density, *B. burgdorferi* reservoir density, small mammal richness, small mammal density, and white-tailed deer density. Models included a random effect of county. All models included covariates of annual mean temperature and annual precipitation. Lyme Disease models included covariates of CDC reporting type. $R^2$ values represent marginal/conditional (fixed/random + fixed) $R^2$.

| Model/Variable | Estimate (SE) | DF | Chisq | **P** |
|---|---|---|---|---|
| Lyme Disease (*w* = 0.95, $R^2$ = 0.17/0.79, *n* = 95, $X^2(5)$ = 41.757, p < 0.001) | | | | |
| Black-legged tick density | -0.405 (0.13) | 1 | 9.27 | 0.002 |
| *B. burgdorferi* reservoir density | 0.243 (0.22) | 1 | 1.19 | 0.276 |
| Small mammal richness | 0.014 (0.08) | 1 | 0.03 | 0.852 |
| Black-legged tick density* *B. burgdorferi* reservoir density | 0.287 (0.06) | 1 | 24.61 | < 0.001 |
| *B. burgdorferi* reservoir density *Small mammal richness | -0.085 (0.03) | 1 | 8.07 | 0.005 |
| Anaplasmosis (*w* = 0.71, $R^2$ = 0.30/0.86, *n* = 116, $X^2(4)$ = 16.935, p = 0.002) | | | | |
| Black-legged tick density | -1.580 (0.61) | 1 | 3.63 | 0.01 |
| Small mammal density | -0.060 (0.04) | 1 | 3.32 | 0.129 |
| Small mammal richness | 0.359 (0.13) | 1 | 6.05 | 0.005 |
| Black-legged tick density*Small mammal density | 0.375 (0.11) | 1 | 8.31 | <0.001 |
| Babesiosis (*w* = 0.50, $R^2$ = 0.73/0.74, *n* = 57, $X^2(4)$ = 16.357, p = 0.003) | | | | |
| Black-legged tick density | 0.240 (0.12) | 1 | 4.31 | 0.038 |
| Small mammal density | 0.875 (0.63) | 1 | 1.94 | 0.164 |
| Small mammal richness | 0.657 (0.55) | 1 | 1.42 | 0.234 |
| Small mammal density *Small mammal richness | -0.120 (0.06) | 1 | 3.36 | 0.067 |
| Ehrlichiosis (*w* = 0.47, $R^2$ = 0.53/0.54, *n* = 116, $X^2(4)$ = 24.886, p < 0.001) | | | | |
| Lone star tick density | 1.533 (0.50) | 1 | 9.30 | 0.002 |
| Deer density | 2.092 (0.54) | 1 | 14.97 | <0.001 |
| Small mammal richness | -0.212 (0.11) | 1 | 3.77 | 0.052 |
| Lone star tick density * Deer density | -0.624 (0.27) | 1 | 5.52 | 0.019 |

Table S6. Regression coefficients and statistics from best-fit models when only sites with forest land cover are included in the data. Predictor variables were blacklegged tick density, lone star tick density, *B. burgdorferi* reservoir density, small mammal richness, small mammal density, and white-tailed deer density. Models included a random effect of county. All models included covariates of annual mean temperature and annual precipitation. Lyme Disease models included covariates of CDC reporting type. $R^2$ values represent marginal/conditional (fixed/random + fixed) $R^2$.

| Model/Variable | Estimate (SE) | DF | Chisq | **P** |
|---|---|---|---|---|
| Lyme Disease ($w$ = 0.99, $R^2$ = 0.24/0.78, $n$ = 81, $X^2(5)$ = 38.584, p < 0.001) | | | | |
| Black-legged tick density | -0.178 (0.13) | 1 | 1.84 | 0.175 |
| *B. burgdorferi* reservoir density | 0.308 (0.22) | 1 | 1.88 | 0.170 |
| Small mammal richness | 0.081 (0.08) | 1 | 0.96 | 0.327 |
| Black-legged tick density* *B. burgdorferi* reservoir density | 0.225 (0.05) | 1 | 20.22 | < 0.001 |
| *B. burgdorferi* reservoir density *Small mammal richness | -0.104 (0.03) | 1 | 11.86 | < 0.001 |
| Anaplasmosis ($w$ = 0.59, $R^2$ = 0.40/0.82, $n$ = 104, $X^2(4)$ = 14.841, p = 0.005) | | | | |
| Black-legged tick density | -0.880 (0.46) | 1 | 3.63 | 0.057 |
| Small mammal density | -0.070 (0.04) | 1 | 3.32 | 0.068 |
| Small mammal richness | 0.299 (0.12) | 1 | 6.05 | 0.014 |
| Black-legged tick density*Small mammal density | 0.209 (0.07) | 1 | 8.31 | 0.004 |
| Babesiosis ($w$ = 0.33, $R^2$ = 0.73/0.74, $n$ = 50, $X^2(4)$ = 16.357, p = 0.003) | | | | |
| Black-legged tick density | 0.202 (0.11) | 1 | 4.31 | 0.068 |
| Small mammal density | 0.686 (0.57) | 1 | 1.94 | 0.231 |
| Small mammal richness | 0.529 (0.50) | 1 | 1.42 | 0.290 |
| Small mammal density *Small mammal richness | -0.099 (0.06) | 1 | 3.36 | 0.098 |
| Ehrlichiosis ($w$ = 0.24, $R^2$ = 0.47/0.49, $n$ = 104, $X^2(3)$ = 20.572, p < 0.001) | | | | |
| Lone star tick density | 0.236 (0.08) | 1 | 7.99 | 0.005 |
| Deer density | 1.570 (0.44) | 1 | 12.56 | < 0.001 |
| Small mammal richness | -0.222 (0.11) | 1 | 4.22 | 0.040 |

Table S7. NEON data products downloaded on 1 November 2019. Note: not all data for all sites was available for the date range requested. See Table S1 for range of years included at each NEON site.

| Data Product ID | Product Name | Site ID | Date Range |
|---|---|---|---|
| DP1.10093.001 | Ticks sampled using drag cloths | All available | 1 Jan 2014 - 31 December 2018 |
| DP1.10092.001 | Tick-borne pathogen status | All available | 2 Jan 2014 - 31 December 2017 |
| DP1.10072.001 | Small mammal box trapping | All available | 3 Jan 2014 - 31 December 2018 |

Table S8. Number of blacklegged (*Ixodes scapularis*) and lone star (*Amblyomma americanum*) ticks tested for tick-borne disease pathogens and number of ticks that tested positive for *Borrelia burgdorferi* (*Bb*), *Anaplasma phagocytophilum* (*Ap*), *Babesia microti* (*Bm*), and *Ehrlichia chaffeensis* (*Ec*) across NEON sites and years of sampling.

| Site | Year | Blacklegged Ticks Tested | *Bb* Pos. | *Ap* Pos. | *Bm* Pos. | Lone star Ticks Tested | *Ec* Pos. |
|---|---|---|---|---|---|---|---|
| BLAN | 2016 | 37 | 7 | 2 | 0 | 3 | 0 |
| BLAN | 2017 | 79 | 23 | 4 | 0 | 2 | 0 |
| DELA | 2015 | --- | --- | --- | --- | 2 | 0 |
| HARV | 2015 | 42 | 16 | 4 | 2 | --- | --- |
| KONA | 2017 | --- | --- | --- | --- | 14 | 0 |
| KONZ | 2015 | --- | --- | --- | --- | 7 | 0 |
| KONZ | 2016 | --- | --- | --- | --- | 122 | 2 |
| KONZ | 2017 | --- | --- | --- | --- | 28 | 0 |
| LENO | 2016 | 1 | 0 | 0 | 0 | 28 | 0 |
| ORNL | 2014 | 4 | 0 | 0 | 0 | 237 | 0 |
| ORNL | 2016 | 23 | 1 | 0 | 0 | 283 | 0 |
| ORNL | 2017 | 3 | 0 | 0 | 0 | 404 | 1 |
| OSBS | 2014 | --- | --- | --- | --- | 209 | 3 |
| OSBS | 2015 | --- | --- | --- | --- | 80 | 0 |
| OSBS | 2016 | --- | --- | --- | --- | 574 | 3 |
| OSBS | 2017 | --- | --- | --- | --- | 320 | 1 |
| SCBI | 2014 | 158 | 10 | 2 | 0 | 37 | 0 |
| SCBI | 2016 | 59 | 14 | 1 | 0 | 11 | 0 |
| SCBI | 2017 | 34 | 6 | 2 | 0 | 80 | 0 |
| SERC | 2016 | 60 | 15 | 0 | 0 | 61 | 0 |
| SERC | 2017 | 82 | 23 | 1 | 0 | 242 | 2 |
| TALL | 2014 | --- | --- | --- | --- | 240 | 0 |
| TALL | 2015 | --- | --- | --- | --- | 187 | 0 |
| TALL | 2016 | --- | --- | --- | --- | 216 | 0 |
| TALL | 2017 | --- | --- | --- | --- | 421 | 0 |
| TREE | 2016 | 117 | 19 | 6 | 4 | --- | --- |
| TREE | 2017 | 112 | 28 | 3 | 7 | --- | --- |
| UKFS | 2015 | --- | --- | --- | --- | 257 | 1 |
| UKFS | 2016 | --- | --- | --- | --- | 328 | 5 |
| UKFS | 2017 | --- | --- | --- | --- | 361 | 0 |

Table S9. List of small mammal species captured across NEON sites.

| Species | NEON Sites |
|---|---|
| *Ammospermophilus harrisii* | SRER |
| *Baiomys taylori* | CLBJ |
| *Blarina brevicauda* | BART, BLAN, GRSM, HARV, KONZ, MLBS, NOGP, ORNL, SCBI, SERC, STEI, TALL, TREE, UNDE, WOOD |
| *Blarina carolinensis* | DELA, DSNY, JERC, LENO, OSBS, TALL |
| *Blarina hylophaga* | KONZ, UKFS |
| *Callospermophilus lateralis* | RMNP, STER |
| *Chaetodipus baileyi* | SRER |
| *Chaetodipus californicus* | SJER |
| *Chaetodipus eremicus* | JORN, SRER |
| *Chaetodipus hispidus* | CLBJ, CPER, KONA, KONZ, OAES, STER |
| *Chaetodipus intermedius* | JORN, SRER |
| *Chaetodipus penicillatus* | JORN, SRER |
| *Cryptotis parva* | BLAN, CLBJ, DSNY, JERC, OSBS, TALL |
| *Dicrostonyx groenlandicus* | TOOL |
| *Didelphis virginiana* | JERC, KONZ, ORNL |
| *Dipodomys merriami* | JORN, SRER |
| *Dipodomys microps* | ONAQ |
| *Dipodomys ordii* | CPER, JORN, MOAB, ONAQ, SRER, STER |
| *Dipodomys spectabilis* | JORN, ONAQ, SRER |
| *Glaucomys sabrinus* | BART, STEI, TREE, UNDE, WREF |
| *Glaucomys volans* | BART, HARV, MLBS, OSBS, SCBI, SERC, STEI, TREE, UNDE |
| *Ictidomys tridecemlineatus* | CPER, DCFS, KONZ, NOGP, OAES, STER, WOOD |
| *Lemmiscus curtatus* | NIWO, ONAQ, RMNP |
| *Lemmus trimucronatus* | BARR, BONA, HEAL |
| *Lepus americanus* | DEJU |
| *Microtus californicus* | SOAP |
| *Microtus longicaudus* | ABBY, CPER, TEAK |
| *Microtus montanus* | NIWO, TEAK |
| *Microtus ochrogaster* | CPER, KONA, KONZ, NOGP, OAES, STER, UKFS |
| *Microtus oeconomus* | BARR, DEJU, TOOL |
| *Microtus oregoni* | ABBY, WREF |
| *Microtus pennsylvanicus* | BLAN, BONA, CPER, DCFS, HARV, HEAL, NOGP, SCBI, STEI, STER, TREE, UNDE, WOOD, YELL |
| *Microtus pinetorum* | BLAN, CLBJ, HARV, JERC, KONZ, MLBS, SCBI, SERC, TALL |
| *Microtus xanthognathus* | BONA |
| *Mus musculus* | BLAN, CLBJ, DSNY, GUAN, JERC, KONA, LAJA, NOGP, OAES, OSBS, SCBI, SERC, STER |
| *Mustela erminea* | ABBY, UNDE, WOOD |
| *Mustela frenata* | HARV, ONAQ, SCBI, STER, UNDE, WOOD |
| *Mustela nivalis* | HARV, NOGP, WOOD |
| *Myodes gapperi* | BART, HARV, MLBS, NIWO, RMNP, STEI, TREE, UNDE, WREF, YELL |
| *Myodes rutilus* | BONA, DEJU, HEAL, TOOL |
| *Napaeozapus insignis* | BART, HARV, MLBS, STEI, TREE, UNDE |
| *Neotoma albigula* | JORN, MOAB, SRER |
| *Neotoma floridana* | DELA, DSNY, JERC, KONZ, LENO, OSBS, TALL, UKFS |
| *Neotoma lepida* | ONAQ |
| *Neotoma mexicana* | MOAB |
| *Neotoma micropus* | CLBJ, JORN, OAES |
| *Neurotrichus gibbsii* | ABBY, WREF |
| *Ochotona princeps* | NIWO |
| *Ochrotomys nuttalli* | GRSM, ORNL, OSBS, TALL |
| *Onychomys arenicola* | JORN |
| *Onychomys leucogaster* | CPER, DCFS, JORN, MOAB, OAES, ONAQ, SRER, STER |
| *Onychomys torridus* | JORN, SRER |
| *Oryzomys palustris* | DELA, DSNY, LENO, OSBS |
| *Perognathus amplus* | JORN, SRER |
| *Perognathus flavescens* | MOAB, OAES, SRER |
| *Perognathus flavus* | CPER, JORN, MOAB, OAES, SRER, STER |
| *Perognathus inornatus* | SJER |
| *Perognathus parvus* | MOAB, ONAQ |
| *Peromyscus attwateri* | CLBJ |
| *Peromyscus boylii* | SJER, SOAP, SRER, TEAK |
| *Peromyscus californicus* | SJER |
| *Peromyscus eremicus* | SRER |
| *Peromyscus gossypinus* | DELA, DSNY, GRSM, JERC, LENO, OSBS, TALL |

Table S9 cont.

| | |
|---|---|
| *Peromyscus keeni* | ABBY, WREF |
| *Peromyscus leucopus* | BART, BLAN, CLBJ, DELA, GRSM, HARV, JORN, KONA, KONZ, LENO, MLBS, NOGP, OAES, ORNL, SCBI, SERC, SRER, STEI, TALL, TREE, UKFS, UNDE |
| *Peromyscus maniculatus* | ABBY, BART, BLAN, CLBJ, CPER, DCFS, GRSM, HARV, KONA, KONZ, MLBS, MOAB, NIWO, NOGP, OAES, ONAQ, ORNL, RMNP, SCBI, SERC, SJER, SOAP, STEI, STER, TEAK, TREE, UKFS, UNDE, WOOD, WREF, YELL |
| *Peromyscus merriami* | SRER |
| *Peromyscus polionotus* | JERC, OSBS, TALL |
| *Peromyscus truei* | MOAB, ONAQ, SJER, SOAP |
| *Phenacomys intermedius* | NIWO |
| *Rattus norvegicus* | NOGP |
| *Rattus rattus* | BLAN, GUAN, LAJA, SCBI |
| *Reithrodontomys fulvescens* | CLBJ, JORN, OAES, SRER |
| *Reithrodontomys humulis* | DSNY, JERC, ORNL |
| *Reithrodontomys megalotis* | CPER, DCFS, JORN, KONA, KONZ, MOAB, NOGP, ONAQ, SJER, SOAP, SRER, STER, UKFS, WOOD |
| *Reithrodontomys montanus* | CLBJ, CPER, OAES, SRER, STER |
| *Sciurus carolinensis* | OSBS, SERC |
| *Sigmodon arizonae* | SRER |
| *Sigmodon hispidus* | CLBJ, DELA, DSNY, JERC, JORN, KONA, KONZ, LENO, OAES, ORNL, OSBS, TALL, UKFS |
| *Sorex arcticus* | STEI, TREE, UNDE |
| *Sorex bairdi* | ABBY, WREF |
| *Sorex cinereus* | BART, BONA, DCFS, DEJU, HARV, HEAL, MLBS, NIWO, NOGP, RMNP, SCBI, STEI, TREE, UNDE, WOOD, YELL |
| *Sorex fumeus* | BART, HARV, MLBS |
| *Sorex haydeni* | DCFS, NOGP, WOOD |
| *Sorex hoyi* | HEAL, TREE, UNDE |
| *Sorex longirostris* | JERC, OSBS |
| *Sorex merriami* | RMNP |
| *Sorex monticolus* | ABBY, NIWO, RMNP, TOOL, WREF |
| *Sorex palustris* | TEAK, UNDE |
| *Sorex trowbridgii* | ABBY, SOAP, TEAK, WREF |
| *Sorex tundrensis* | BONA, HEAL |
| *Sorex ugyunak* | TOOL |
| *Sorex vagrans* | ABBY, WREF |
| *Spermophilus armatus* | YELL |
| *Spermophilus franklinii* | WOOD |
| *Spermophilus parryii* | TOOL |
| *Spermophilus spilosoma* | CPER, JORN, MOAB |
| *Spermophilus tereticaudus* | SRER, STER |
| *Sylvilagus audubonii* | JORN, MOAB, SRER |
| *Sylvilagus floridanus* | CLBJ, CPER, KONA, KONZ, SCBI |
| *Sylvilagus nuttallii* | MOAB, RMNP |
| *Synaptomys cooperi* | TREE, UNDE |
| *Tamias amoenus* | YELL |
| *Tamias dorsalis* | ONAQ |
| *Tamias minimus* | MOAB, NIWO, ONAQ, RMNP, UNDE, YELL |
| *Tamias quadrivittatus* | RMNP |
| *Tamias rufus* | MOAB |
| *Tamias speciosus* | TEAK |
| *Tamias striatus* | BART, BLAN, GRSM, HARV, MLBS, ORNL, SCBI, SERC, STEI, TALL, TREE, UKFS, UNDE |
| *Tamias townsendii* | ABBY, WREF |
| *Tamiasciurus douglasii* | ABBY, SJER |
| *Tamiasciurus hudsonicus* | BART, BONA, DEJU, HARV, HEAL, NIWO, RMNP, TREE, UNDE |
| *Thomomys talpoides* | WOOD, YELL |
| *Zapus hudsonius* | BLAN, BONA, DCFS, HARV, KONZ, MLBS, NOGP, SCBI, SERC, STEI, TREE, UNDE, WOOD |
| *Zapus princeps* | DCFS, NIWO, NOGP, WOOD, YELL |
| *Zapus trinotatus* | ABBY |

Figure S4. Correlation matrix of predictor variables in generalized linear effects models (GLMMs; Table SX). Absolute values of correlations >0.6 are highlighted in yellow. Only 3 pairs of variables indicate high levels of correlation: temperature and latitude, density of all blacklegged ticks (*Ixodes scapularis* and *Ixodes pacificus*) and density of nymphal blacklegged ticks (*I. scapularis* and *I. pacificus*), and density of all lone star ticks (*Amblyomma americanum*) and density of nymphal lone star ticks (*A. americanum*). Of these pairs of variables, only temperature and latitude jointly appear in GLMMs. Despite the high levels of correlation, the inclusion of both variables in the GLMMs is important to capture known ecological/behavioral gradients of blacklegged ticks along each variable.

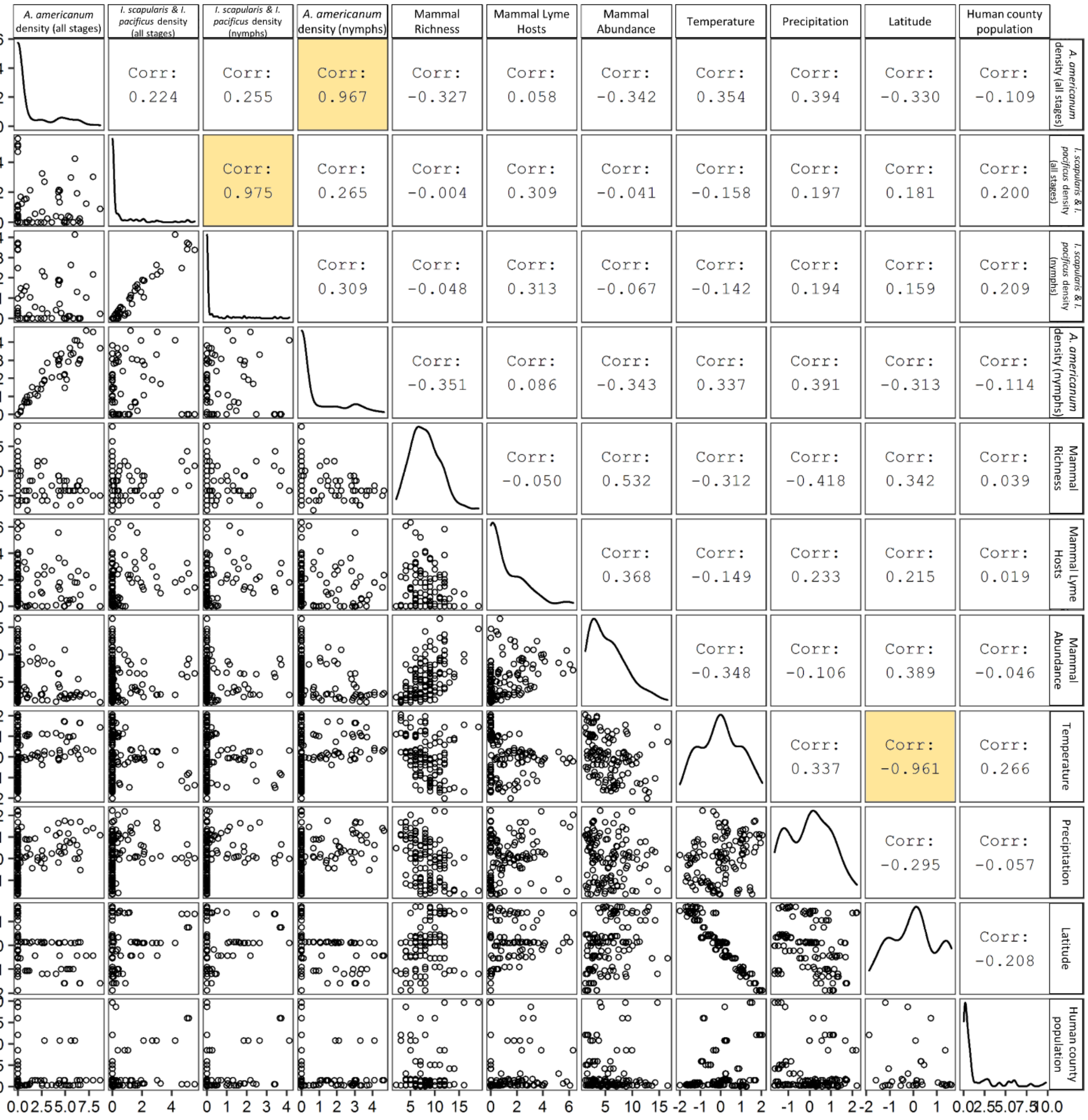
